# Non-Synonymous, Synonymous, and Non-Coding Nucleotide Variants Contribute to Recurrently Altered Biological Processes During Retinoblastoma Progression

**DOI:** 10.1101/2022.10.27.512289

**Authors:** Kevin Stachelek, Narine Harutyunyan, Susan Lee, Assaf Beck, Jonathan Kim, Liya Xu, Jesse L. Berry, Aaron Nagiel, C. Patrick Reynolds, A. Linn Murphree, Thomas C. Lee, Jennifer G. Aparicio, David Cobrinik

## Abstract

Retinoblastomas form in response to biallelic *RB1* mutations or *MYCN* amplification and progress to more aggressive and therapy-resistant phenotypes through accumulation of secondary genomic changes. Progression-related changes include recurrent somatic copy number alterations and typically non-recurrent nucleotide variants, including synonymous and non-coding variants, whose significance has been unclear. To assess synonymous and non-coding variant contributions to recurrently altered processes, we identified altered genes and over-represented variant gene ontologies in 168 exome or whole-genome-sequenced retinoblastomas and 12 tumor-matched cell lines. In addition to initiating *RB1* mutations, *MYCN* amplification, and established retinoblastoma SCNAs, the analyses revealed enrichment of variant genes related to diverse biological processes including histone monoubiquitination, mRNA processing (P) body assembly, and mitotic sister chromatid segregation and cytokinesis. Importantly, inclusion of non-coding and synonymous variants increased the enrichment significance of each over-represented biological process term. To assess the effects of such mutations, we performed functional tests of 3’ UTR variants of *PCGF3* (a BCOR-binding component of Polycomb repressive complex I) and *CDC14B* (a regulator of sister chromatid segregation) and a synonymous variant of *DYNC1H1* (a regulator of P-body assembly). *PCGF3* and *CDC14B* 3’ UTR variants impaired gene expression whereas a base-edited *DYNC1H1* synonymous variant altered protein structure and stability. Compared to tumors, retinoblastoma cell lines had a partially overlapping variant gene spectrum and enrichment for p53 pathway mutations. These findings reveal potentially important differences in retinoblastoma cell lines and antecedent tumors and implicate synonymous and non-coding variants, along with non-synonymous variants, in retinoblastoma oncogenesis.

## Introduction

Cancers develop in response to successive genetic or epigenetic changes in a susceptible cell-of-origin and progress to more aggressive phenotypes via continuous mutation and selection. Whereas secondary genomic changes may be present in only a subset of tumor cells, subclonal mutations can increase tumor aggressiveness and therapy resistance ^1^. However, mechanisms underlying tumor progression are poorly defined, in part due to challenges identifying recurrently altered progression driver genes and cellular functions.

Retinoblastoma is a pediatric retinal cancer that may be used to dissect the role of progression-related subclonal mutations, as most retinoblastomas initiate in response to the same genomic alteration, pass through similar pre-malignant stages, and carry low mutational burdens. ∼98% of retinoblastomas form in response to biallelic inactivation of *RB1* and are thought to originate in maturing cone precursors and proceed through a premalignant state before retinoblastomas appear, whereas ∼ 2% form in immature cone precursors in response to *MYCN* amplification ^2–5^. *RB1*-mutant retinoblastomas may initially lack clonal chromosomal abnormalities or single nucleotide variants (SNVs) in exomic sequences of protein coding genes beyond *RB1* ^6^, implying that progression-related genomic changes are superimposed over otherwise clean genomic landscapes.

Most *RB1*-mutant retinoblastomas that are sufficiently advanced to require enucleation have one or more of the recurrent somatic copy number alterations (SCNAs) 1q+, 2p+, 6p+,7q+, 16q-, and 19q+ ^7^. In most instances, these ‘retinoblastoma SCNAs’ are likely acquired after retinoblastomas emerge, as they are usually present at subclonal frequencies, are more common in larger and more advanced retinoblastomas, and may appear during retinoblastoma therapy ^7–9^. While some SCNAs have prognostic value (8), their progression-related cell signaling effects are unknown. In contrast to highly recurrent SCNAs, prior whole genome or exome sequencing (WGS/WES) of 156 treatment-naïve retinoblastomas revealed no significantly recurrently altered genes with exome tier-1 mutations, according to standard metrics ^10^, despite identification of *BCOR* variants in 20, *CREBBP* variants in three, and 10 genes in two samples (Supplementary Tables S1, S2) ^6,11–13^. A similar number of genes had multiple mutations in a more recent analysis of 103 retinoblastomas ^14^ and ultra-sensitive targeted sequencing revealed additional mutated genes ^15,16^, yet significantly recurrently altered cellular functions have not been identified.

Gene ontology and protein interaction network analyses may be used to identify recurrently altered cellular functions or provide clues to affected pathways in the absence of significantly recurrently altered genes ^17^. While these approaches are often applied solely to non-synonymous variants with known oncogenic effects or to all non-synonymous variants in a tumor series, such analyses rarely include synonymous and non-coding variants owing to their presumed passenger status. However, the low retinoblastoma somatic variant rate of 0.053 mutations per coding megabase ^18^ suggests that a high proportion of variants –including synonymous and non-coding variants – may drive cancer progression. When reported, synonymous and non-coding variants made up 55% of somatic variants (Supplementary Table S2) ^11–13^, representing a potentially impactful cohort.

Here, we aimed to identify biological processes that are recurrently altered during retinoblastoma progression and to assess the importance of synonymous and non-coding variants in over-represented biological process gene ontologies. We further examined the retention of subclonal progression-related variants during cell line establishment to discern cell line utility in cancer progression studies. Our analyses reveal previously unrecognized recurrently altered cellular functions, surprisingly frequent contributions of synonymous and non-coding mutations to recurrently altered biological process ontologies, and deleterious effects of synonymous and non-coding mutations that contribute to recurrently altered cellular functions, implicating such changes in retinoblastoma progression.

## Materials and Methods

### Tumor and normal tissue

Retinoblastoma tumors and normal tissues were obtained under protocols approved by the Children’s Hospital Los Angeles and Children’s Oncology Group (COG) institutional review boards. Following enucleation, tumors were divided for flash-freezing in liquid nitrogen or cell culture. For the CHLA-VC-RB series, matched normal DNAs were obtained from orbital fat pad adipose-derived mesenchymal cells or buccal swabs (Oragene DISCOVER OGR-575 DNA Collection Kit (DNA Genotek).

### Establishment and propagation of CHLA-VC-RB and CHLA-RB cell lines

CHLA-VC-RB tumors were explanted and cultured in IMDM, 10% fetal bovine serum (FBS), Insulin-Transferrin-Selenium (ITS, Thermo Fisher Scientific), and 4 mM L-glutamine for up to two months, frozen in Cell Freezing Medium-DMSO (Sigma-Aldrich), and propagated by plating without dissociation at ∼ 1-5 × 10^6^ cells per well of a 96-well dish in 200 μl RB culture medium (IMDM (Mediatech), 10% FBS, 2 mM glutamine (Mediatech), 55 μΜ beta-mercaptoethanol (Sigma-Aldrich), and 10 μg/ml insulin (Humulin-R, Lilly)) at 37° C and 5% CO2 ^5^. Regardless of well-size, cells were cultured under ∼ 1 cm media and two-thirds of the media (∼133 μl in the 96-well format) was changed approximately every two days, when media approached amber color (∼pH 7.0). For optimal growth, undissociated cell clusters were maintained at densities sufficient to acidify the media to ∼ pH 7.0 within ∼ 36 h and were passaged as needed to prevent acidification past a toxic pH 6.5 by splitting by no more than 2:3. Cell growth was monitored in terms of 96-well equivalents and days in culture. 1×10^5^ CHLA-VC-RB-29 cells were engrafted subretinally in a NSG mouse as described ^5^ and a tumor appearing at 130 days post-engraftment re-explanted as for primary tumors. CHLA-RB tumors were explanted into IMDM, 20% fetal bovine serum, ITS, and 4 mM L-glutamine and media changed every 1-2 days. After establishment, cell lines were passaged 1:2 every ∼ 10 days. Short tandem repeat (STR) profiles were recorded for all cell lines using the DNA samples used for exome analyses (Supplementary Table S3).

### Exome sequencing and variant calling

DNA was isolated using QIAamp DNA Mini Kit (Qiagen) according to manufacturer’s instructions. Exome sequencing was performed by Macrogen Inc (Seoul, South Korea) or BGI (Shenzhen, China) with SureSelect V5 Exome Capture Kit (Agilent) and sequencing on Illumina HiSeq. Sequences were aligned to UCSC hg19 human reference genome using the Burrows-Wheeler aligner bwa-mem http://bio-bwa.sourceforge.net/. Adapters were trimmed with trimmomatic ^19^, duplicate reads removed with Picard Markduplicates http://broadinstitute.github.io/picard/, and base quality scores recalibrated with GATK ^20^. Coverage depth was determined with Mosdepth based on SureSelect V5 capture regions ^21^.

Somatic variants in protein-coding genes were called using GATK MuTect2 ^22^ and Strelka2 ^23^ in CHLA-VC-RB tumors and cell lines relative to matched normal DNAs and using MuTect2 in tumor-only mode in CHLA-RB cell lines. Systematic sequencing artifacts were identified using the 12 CHLA-VC-RB matched normal DNAs sequenced in parallel. Somatic variants not present in the matched or panel of normal DNAs were annotated using Ensembl Variant Effect Predictor ^24^. Annotated variants were filtered using the VariantAnnotation R package from Bioconductor ^25^. Variants were removed if documented in the Genome Aggregation Database (gnomAD) ^26^ at an allele frequency > 0.05% in sequenced human genomes ^27^. CHLA-RB cell line variants were also excluded if annotated in dbSNP. Variants were further filtered if called at variant allele frequency (VAF) < 0.05 or read depth of < 3 but were reported when VAF < 0.05 and detected solely in a sample and a matched tumor or cell line. To identify variants in matched samples that were not detected by variant calling algorithms, the allele frequency of each called variant was calculated in all other samples using samtools mpileup and bcftools with minimal qc thresholds. Variants detected in RB-VC-CHLA24-T and at higher levels in RB-VC-CHLA46-CL were deemed to represent contamination and removed. Code used for variant annotation, filtration, and gene ontology analysis is available as an R package and workflow at https://github.com/cobriniklab/stchlkExome and https://github.com/cobriniklab/rb_exome.

### Somatic copy number alterations (SCNAs)

SCNAs were detected using the Bioconductor R package CopywriteR which uses off-target exome sequencing reads to define a uniformly distributed copy number profile by circular binary segmentation ^28^. Read density was measured with a bin size of 50 kb. Exomes of matched normal DNAs (from orbital fat pad-derived cells or buccal swabs) were used to determine relative copy number for CHLA-VC-RB tumors and cell lines whereas absolute copy number was measured for CHLA-RB cell lines.

### Allelic Imbalance

All somatic variant sites detected by MuTect2 were collected into a single .bed file. B allele frequencies for all samples were calculated using the mpileup2baf Perl script https://github.com/cobriniklab/rb_exome/blob/master/LOH/mpileup2baf.pl shared by Irsan Kooi (VU University). B allele frequency segmentation was performed with BAF segmentation using 50 kb non-overlapping bins ^29^, yielding a profile of mirrored B allele frequency (mBAF) representing absolute deviation from heterozygous allele frequency over a given binned region. A region was designated as having allelic imbalance if mBAF was greater than 0.56 ^29^. A region was designated as having loss of heterozygosity (LOH) if mBAF was greater than 0.7. *RB1* allelic imbalance is reported for segments spanning hg19 chr13 48877887 - 49056122.

### Gene ontology analysis

Functional enrichment analysis of somatic variants in Supplementary Table S1 was performed with the WebGestaltR R package ^30^. Over-representation of gene symbols present in Gene Ontology biological processes (release 2021-02, at http://geneontology.org/docs/ontology-documentation/) was quantified by a hypergeometric test. For terms initially having FDR ≤ 0.5, significance tests were amended to reflect multiple instances of repeatedly mutated genes. Ontology terms with a false discovery rate (FDR) adjusted by Benjamini-Hochberg correction are reported.

### Luciferase reporter assay

3’ UTR sequences of *PCGF3* and *CDC14B* were amplified from H9 embryonic stem cells using primers *PCGF3*f, *PCGF3*r, *CDC14B*f, and *CDC14B*r (Table S4) and inserted in place of SV40 UTR and poly (A) sequences in pGL3-SV40 using in-Fusion (Takara). Mutations were generated in 3’ UTR sequences using In-Fusion and mutagenic primers *PCGF3*-inf-f, *PCGF3*-inf-r, *CDC14B*-inf-1-f, *CDC14B*-inf-1-r, *CDC14B*-inf-2-f, and *CDC14B*-inf-2-r; mutated positions underlined (Table S4). The resulting plasmids were transfected into Y79 retinoblastoma cells with 5 μg of each plasmid, 2 × 10^5^ cells in 100 μl, with 2 pulses of 20 ms at 1100 V using the Neon transfection system (Thermo Fisher Scientific). After 48 h incubation, luminescence was measured by Nano-Glo® Dual-Luciferase® Reporter Assay (Promega).

### CRISPR base editing

We identified a gRNA and cytidine base editor to generate the chr14:102,442,053 *DYNC1H1* c.261C >T mutation using a custom script that applies BE-designer (http://www.rgenome.net/be-designer/) to diverse base editors (https://github.com/cobriniklab/gRNA-for-BE-design-repository). The identified gRNA coding sequence (gagga<u>C</u>gtcggtgatgaagg; targeted C underlined) was inserted into pSPgRNA (Addgene plasmid # 47108). CHLA-VC-RB31 cells were co-transfected with the resulting pSPgRNA-*DYNC1H1-mut* or a control pSPgRNA expressing non-targeting gRNA and dCas9-cytosine base editor vector pCAG-CBE4max-SpG-P2A-EGFP (RTW4552) (Addgene plasmid # 139998) ^31^ using the Neon transfection system with 5 μg of each plasmid, 2 × 10^5^ cells in 100 μl tips, and transfected with 2 pulses of 20 ms at 1100 V. After 24 hours culture, transfected GFP+ cells were sorted to yield bulk GFP+ cultures and single cell clones. Editing was assessed by PCR of the targeted region using primers *DYNC1H1-1f* and *DYNC1H1-ex2r* followed by Sanger sequencing. After identification of clones with heterozygous edits a second round of transfection yielded homozygous edited clones with no local off-target edits confirmed by Sanger sequencing.

### RT-qPCR

RNA was extracted from base-edited and control CHLA-VC-RB31 cells using miRNeasy Kit (Qiagen); cDNA synthesized using iScript cDNA Synthesis Kit (BioRad); RT-qPCR performed with iTaq Universal SYBR Green Supermix (BioRad) and exon-specific primers (Supplementary Table S4). Fold change was calculated using the ∆∆Ct method with *GAPDH* as reference and values normalized to the mean of the nontargeted sample.

### Immunoblotting and protein stability analyses

Cell lysates were prepared by suspending 1–2 × 10^5^ cells in lysis buffer (RIPA (MilliporeSigma), EDTA-free Protease Inhibitor Cocktail (cOmplete, Roche), 0.1% SDS) on ice for 30 min and centrifuged at 14,000 rpm for 10 min at 4°C. Lysates were adjusted to 40 μg protein / 12 μl RIPA buffer, treated with papain (Worthington, LK003176) for 30 min at 37° C, combined with 4 μl 4x sample loading buffer (Thermo Fisher), and incubated at 95° C for 5 min. Proteins were separated by electrophoresis through 3-8% gradient SDS polyacrylamide tris-acetate gels (ThermoFisher), and transferred to polyvinylidene fluoride membranes (Cytiva, 10600023). Following transfer, blots were blocked in Tris buffered saline (TBS), 0.1% Tween-20, with 5% nonfat dry milk for 1 hour at room temperature and separately incubated with primary antibodies to DYNC1H1 (Proteintech, 12345-1-AP) at 1:500 and GAPDH (Thermo Fisher, 39-8600) at 1:1000 overnight at 4°C, followed by 3 washes in TBS, 0.1% Tween-20 and incubation with horseradish peroxidase-conjugated secondary antibodies (Santa Cruz) for 1 hour at room temperature.

## Results

### Retinoblastoma samples, exome sequencing, and tumor-initiating mutations

To assess somatic mutations in retinoblastomas and tumor-derived cell lines, we performed exome sequencing on 12 treatment-naïve retinoblastoma tumors, their matched cell lines, and matched normal DNAs and on 15 retinoblastoma cell lines without matched normal DNAs (Supplementary Table S5). Tumor exomes were sequenced to post-processing mean read depths of 168x, 95x, and 99x for tumor, cell line, and normal samples, respectively (Supplementary Fig. S1), ∼3 - 6-fold greater depth than prior studies (Supplementary Table S2). Tumor and cell line SCNAs, allelic imbalance, and loss of heterozygosity (LOH) were identified (Supplementary Tables S6, S7) and somatic variants recorded when VAF > 0.05 and the variant was not detected in matched normal or in 33 unmatched samples. Tumor variants were also recorded when VAF <0.05 and the variant was also detected in a matched cell line (Supplementary Table S8).

To assess the sensitivity of the exome analyses, we evaluated *RB1* and *MYCN* status in all samples. Biallelic *RB1* inactivation via SNVs, indel, deletions, and copy-neutral LOH (CN-LOH) was detected in 9 of 12 CHLA-VC tumor - cell line pairs (Table 1). The three exceptions included a tumor with wild type *RB1* and *MYCN* amplification (CHLA-VC-RB24), a tumor with clinically reported *RB1* promoter methylation and chromosome 13q CN-LOH (CHLA-VC-RB20), and a tumor from a patient with bilateral retinoblastoma and 13q CN-LOH suggestive of a germline promoter or deep intron *RB1* mutation (CHLA-VC-RB41). Biallelic *RB1* inactivation or *MYCN* amplification was also detected in 14 of 15 CHLA-RB cell lines, with the exception (CHLA-RB292) having monoallelic deletion (Table 1). Thus, biallelic *RB1* loss or *MYCN* amplification was detected in 25 of 27 samples.

**Table 1.**
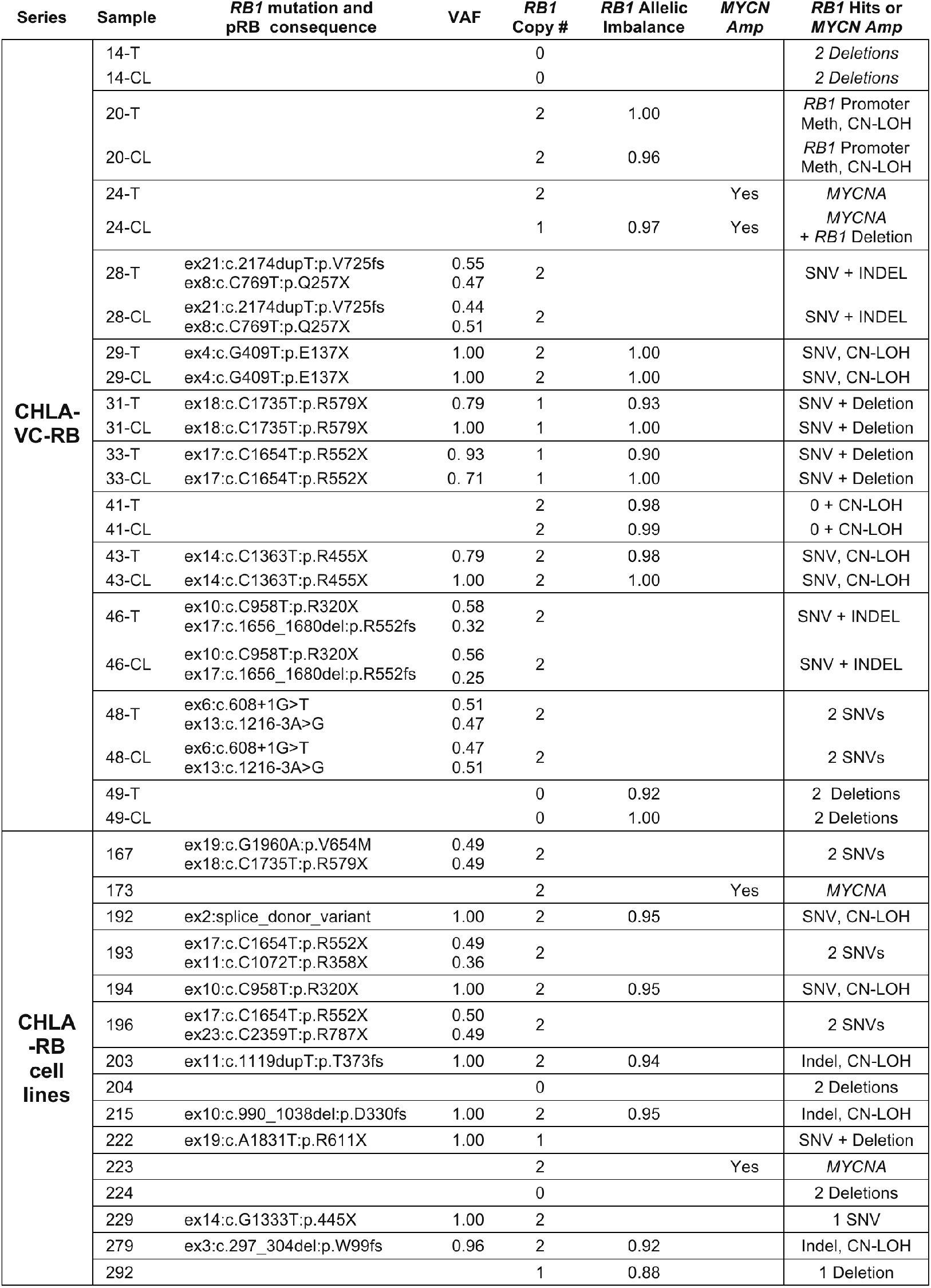
*RB1* and *MYCN* status in retinoblastoma tumors (T) and cell lines (CL). VAF, variant allele frequency. *RB1* Allelic Imbalance, shown if mirrored B allele frequency (mBAF) spanning *RB1* is ≥ 0.56. CN-LOH, copy-neutral loss of heterozygosity, defined as *RB1* mBAF > 0.7 and no *RB1* copy number alteration.

### Somatic variants and repeatedly mutated genes in retinoblastoma tumors

Past WES/WGS of 156 treatment-naïve retinoblastomas identified secondary mutations or focal deletion in *BCOR* (in 20) and *CREBBP* (in 3), and 470 additional somatic mutations, with VAFs from 0.026 to 1.0 (Supplementary Table S1, Fig. 1A). Here, WES of 12 tumors from patients with similar demographics (Supplementary Table S5) revealed 50 SNVs with VAF > 0.05 plus 10 SNVs initially called in cell lines but also present in matched tumors, yielding 60 somatic mutations (mean 5; median 4) (Fig. 1B). The distribution of VAFs was lower than in prior studies, likely reflecting deeper sequencing and retention of variants initially called in cell lines. Among the 60 variants, six affected genes altered in previously sequenced retinoblastomas, including two in *BCOR* and one in *PAN2, NAF1, NCAM1*, and *SAMD9* (Supplementary Table S1). Overall, 14 genes were mutated in at least two samples, of which *BCOR, MYCN*, and *ARID1A* were also repeatedly altered in targeted cancer gene sequencing ^15,16^ (Fig. 1C-E, Supplementary Table S1). However, no genes showed recurrence at rates exceeding a significance threshold of p<0.05 when assessed by MutSig2CV, likely due to the low retinoblastoma somatic mutation rate ^10^.

**Figure 1.**
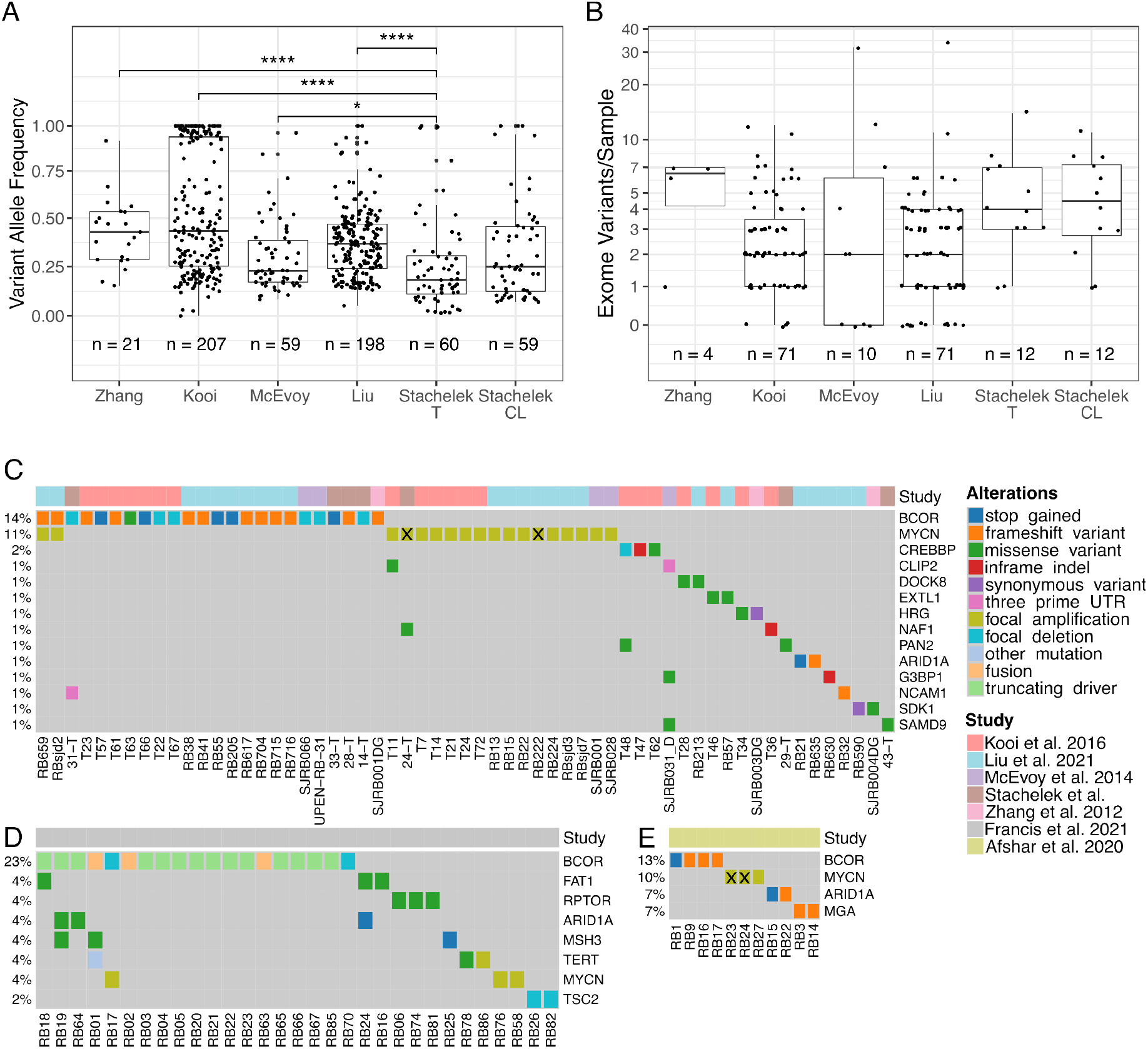
Somatic variant frequencies and repeatedly mutated genes beyond *RB1* in 168 fully reported retinoblastoma tumors. A. Variant allele frequencies (VAF) of all exomic non-*RB1* single nucleotide and indel variants in this and two prior studies ^12,13^, all synonymous and non-synonymous coding sequence variants ^11^, or all non-synonymous coding sequence variants ^6^. B. Number of variants detected per sample in each study displayed with pseudo log scale. Box plot horizontal lines represent median. Differences between the current and prior groups of primary tumors were evaluated with 1-way ANOVA (p = 3.57e-12 (A); p = 0.0698 (B)) and differences between the current tumors and cell lines were evaluated by pair-wise t-tests. All significant differences are indicated. *, p<0.05; ***, p<0.001; ****, p<0.0001. C-E. Repeatedly mutated genes beyond *RB1* identified in retinoblastoma whole genome or whole exome sequencing (C) or in targeted sequencing (D, E). Tumors with focal *MYCN* amplification and no annotated RB1 lesion are indicated with x.

### Recurrently altered cellular functions

Given the rarity of somatic mutations, we hypothesized that a significant proportion affect retinoblastoma progression-related functions. Thus, we examined whether secondary mutations are enriched for specific biological processes gene ontology terms using WebGestalt over-representation analyses ^30^. Our analyses included all somatic SNVs and indels beyond *RB1* identified in this and four prior WES/WGS studies ^6,11–13^ and were performed separately for the 417 variants that altered protein amino acid sequences and for all 542 SNV or indel variants including 37 UTR and 88 synonymous sequences. To mitigate effects of frequently altered genes such as *BCOR*, over-represented ontologies (FDR < 0.25) were identified by inputting each gene name once and statistical significance determined by counting all variants separately. The over-representation analysis did not include variants reported in targeted retinoblastoma sequencing studies ^15,16^, which may omit as-yet-unidentified progression driver genes.

Among ontologies with FDR < 0.25 and comprised of least five variants, the analyses revealed significant over-representation of terms related to *histone monoubiquitination*, c*ovalent chromatin modification, cytoplasmic mRNA processing (P) body assembly, ribonucleoprotein (RNP) complex assembly, integrin-mediated cell adhesion, semaphorin-plexin signaling, mitotic cytokinesis, mitotic sister chromatid segregation*, and *phosphatidylinositol phosphorylation* (Table 2). We also detected over-representation of highly related ontologies with higher (less significant) FDRs and ontologies that lacked specific signaling features, such as *regulation of organelle assembly* and *multi-organism cellular process* (Supplementary Table S9), which were not further considered. Notably, the identified ontologies were largely comprised of genes that were not included in the targeted sequencing panels previously applied to retinoblastoma (Table 2).

**Table 2.**
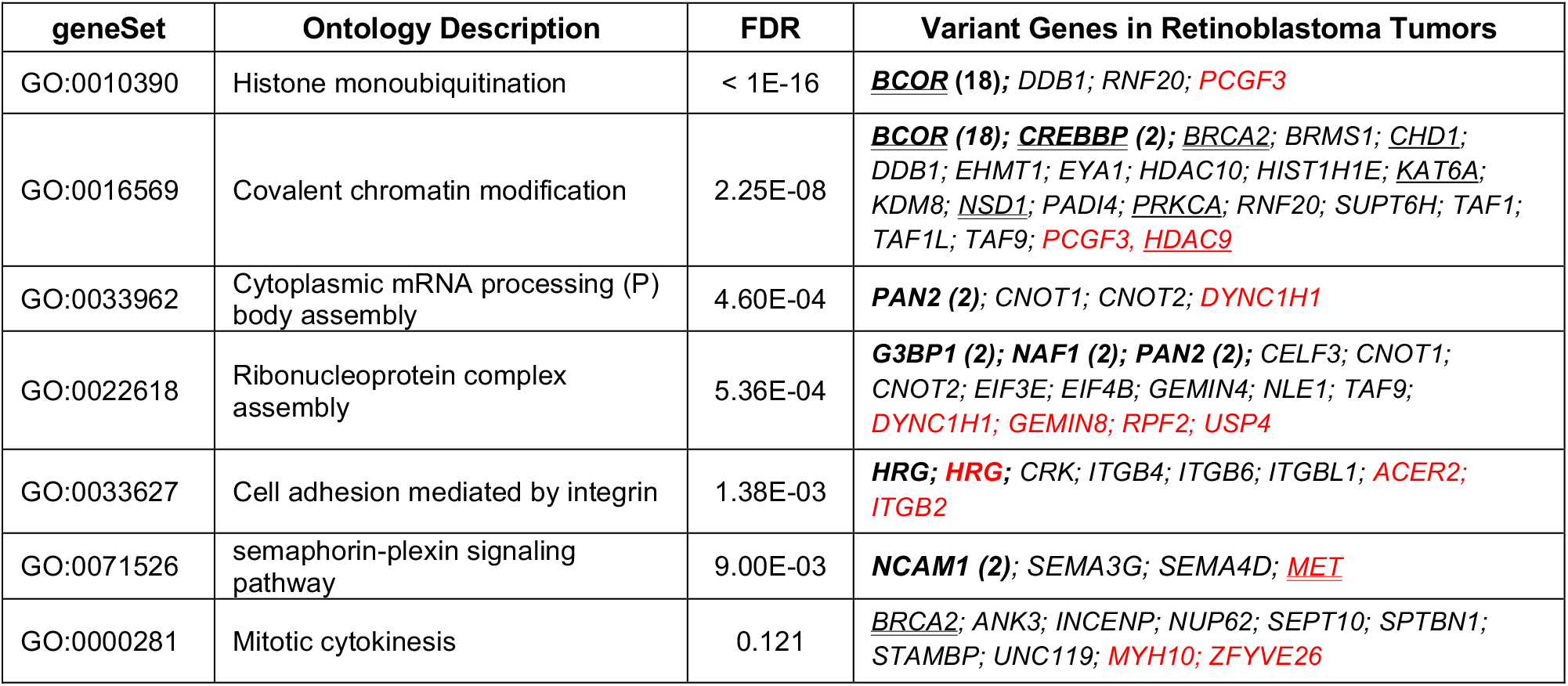
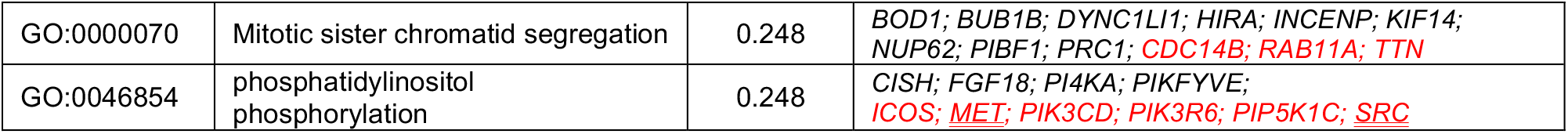
Biological process ontologies over-represented among somatically altered genes in 168 sequenced retinoblastoma tumors. **Bold**, recurrently altered (number of recurrences). Red, non-protein-altering variant. <u>Underlined</u>, genes included in UCSF500. Double underlined, genes included in UCSF500 and MSK-IMPACT versions used for retinoblastoma targeted sequencing. FDR, false discovery rate.

The most significantly over-represented ontology was *histone monoubiquitination*, represented by non-synonymous variants of *BCOR, DDB1, RNF20*, and a non-coding variant of *PCGF3. Covalent chromatin modification* was the next most significant, driven by *BCOR* as well as genes encoding other histone modification writers (*CHD1, CREBBP, DDB1, EHMT1, KAT6A, NSD1, PRKCA, PADI4, PCGF3, RNF20*), erasers (*BRMS1, EYA1, HDAC9, HDAC10, KDM8*), and histones themselves (*HIST1H1E*). Enrichment of covalent chromatin modification was mainly driven by histone monoubiquitination variants, as significance was greatly reduced when monoubiquitination-related genes were omitted (FDR=0.143), and no other chromatin modification category was significantly enriched.

Genes related to RNP assembly fell into several sub-categories, of which *cytoplasmic mRNA P body assembly* was independently over-represented by variants in *PAN2, CNOT1, CNOT2*, and *DYNC1H1* (FDR=4.60E-04). Although *RNP assembly* remained over-represented (FDR = 2.77E-02) when P body assembly genes were removed, other RNPs affected by multiple variants (the SMN splicing/spliceosome complex (*GEMIN4* and *GEMIN8*) ^32^, the H/ACA snRNPs that assemble TERC RNA – TERT complexes (*NAF1*) ^33^, and translation initiation complexes (*EIF4B* and *EIF3E*)) related to dissimilar and not over-represented processes. The other over-represented ontologies were largely distinct from one another, as *mitotic cytokinesis* and *mitotic sister chromatid segregation* had only two variant genes in common (*NUP62* and *INCENP*), *semaphorin-plexin signaling* and *phosphatidylinositol phosphorylation* shared only one gene *(MET*), and *cell adhesion mediated by integrin* shared no variant genes with the other categories. Thus, the analyses revealed over-representation of seven largely distinct processes: histone monoubiquitination, P-body assembly, integrin-mediated cell adhesion, semaphorin-plexin signaling, sister chromatid segregation, cytokinesis, and phosphatidylinositol phosphorylation.

### Contributions of synonymous and non-coding variants to recurrently altered cellular functions

Notably, each of the enriched biological processes had contributions of noncoding UTR or synonymous variants along with non-synonymous variants (Table 2, genes shown in red). Moreover, the significance of six of the seven over-represented biological process terms increased (FDRs decreased) when the 125 non-protein-altering variants were added to the 417 protein-altering variants, whereas *histone monoubiquitination* remained at FDR < 10^−16^ (Fig. 2A). The FDRs are unlikely to have been lowered by chance, as addition of 125 randomly selected genes to the original 417 non-synonymous variants failed to improve the FDR of the seven ontology terms in 68 of 70 iterations (Table S10) (p<0.0001, Fisher’s exact test). Overall, 52 of 417 (12.5%) protein-altering variants and a similar 16 of 125 (12.8%) non-protein-altering variants fell in at least one of these seven enriched biological function categories (p = 0.88, Fisher exact test).

**Figure 2.**
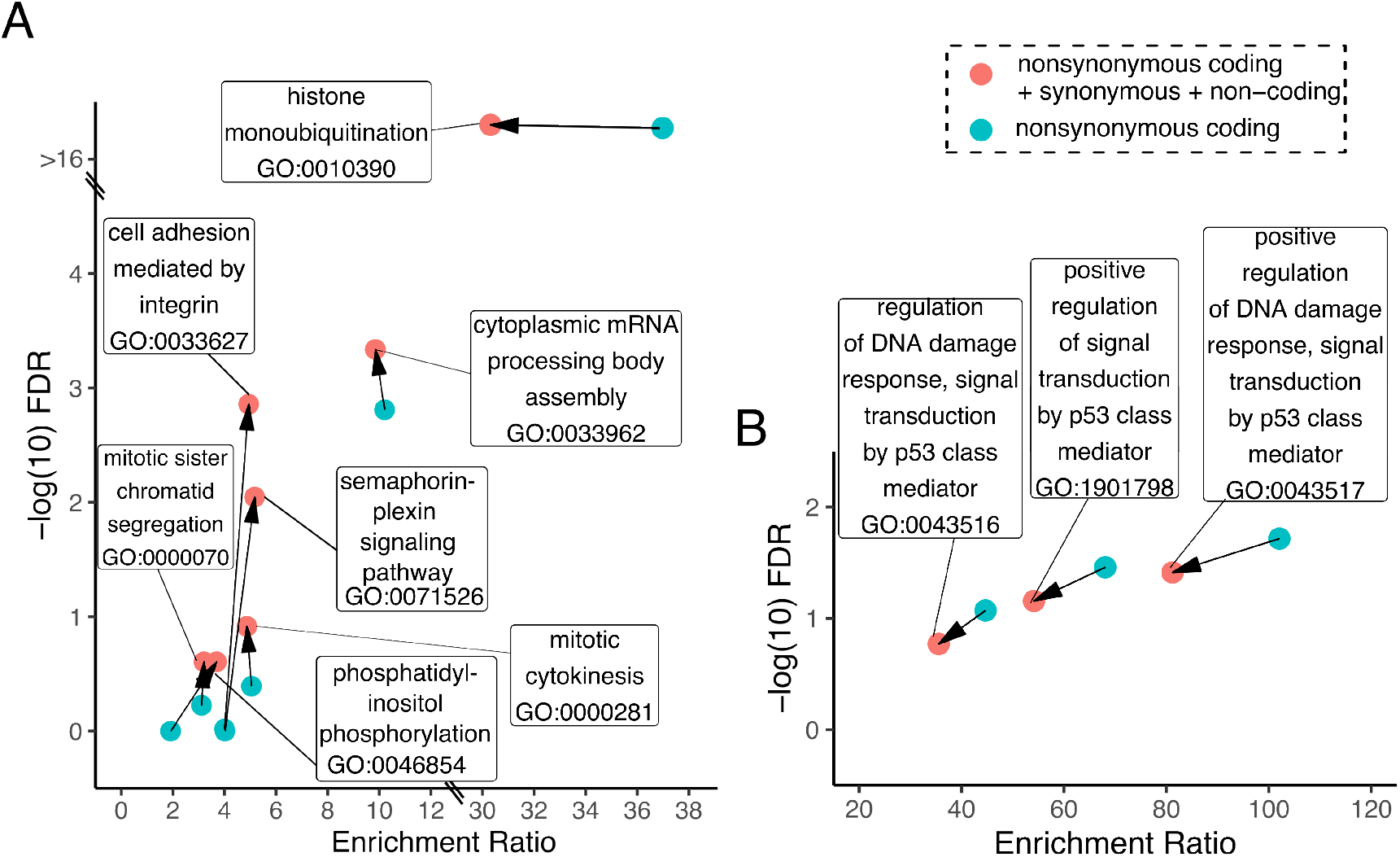
Biological process gene ontology terms over-represented in exome variants in retinoblastoma whole exome or whole genome sequencing. Over-representation analyses were performed separately for non-synonymous variants affecting protein amino acid sequence (blue) and for all exomic variants including synonymous coding mutations and non-coding 5’ or 3’ UTR mutations (red) in 160 treatment-naïve tumors (A) or the 12 CHLA-VC-RB cell lines (B). Arrows indicate effect of non-protein-altering mutations on enrichment ratio and FDR values.

As non-coding and synonymous coding variants contributed to multiple over-represented biological process ontologies and may alter gene and protein expression in a manner that promotes oncogenesis ^34,35^, we explored the effects of such mutations in genes affecting three of the enriched biological processes. We focused on a 3’ UTR variant in *PCGF3* (c.*3764G>A), which contributes to *histone monoubiquitination*, two co-occurring 3’ UTR variants in *CDC14B* (m1=c.*2060G>A; m2=c.*2447A>G,), which contributes to *mitotic sister chromatid segregation*, and a synonymous variant in exon 2 of *DYNC1H1* (c.261C>T), which contributes to *cytoplasmic mRNA P body assembly*.

We hypothesized that 3’ UTR variants affecting *PCGF3* and *CDC14B* might impact RNA stability or translation. In keeping with this possibility, we verified that their variants were within 3’ UTR regions that are expressed in retinoblastoma tumors and that the *PCGF3* variant and *CDC14B* variant m2 are moderately evolutionarily conserved (Supplementary Fig. S2). We then assayed the variant effects on gene expression using pGL3-SV40 luciferase reporter plasmids with the SV40 poly (A) signal replaced by the wild type or mutant *PCGF3* and *CDC14B* 3’ UTRs and inferred poly (A) signals (Fig. 3A). Luciferase assays revealed a significant ∼ 3-fold reduction in luminescence of the mutant versus wild type *PCGF3* UTR in Y79 cells (Fig. 3B). Similarly, the *CDC14B* m1 variant reduced luminescence whereas the m2 variant did not significantly differ from wild type and the combined m1/m2 mutant reversed the *CDC14B* m1 effect (Fig. 3B), behavior consistent with sequential impairment and restoration of *CDC14B* expression.

**Figure 3.**
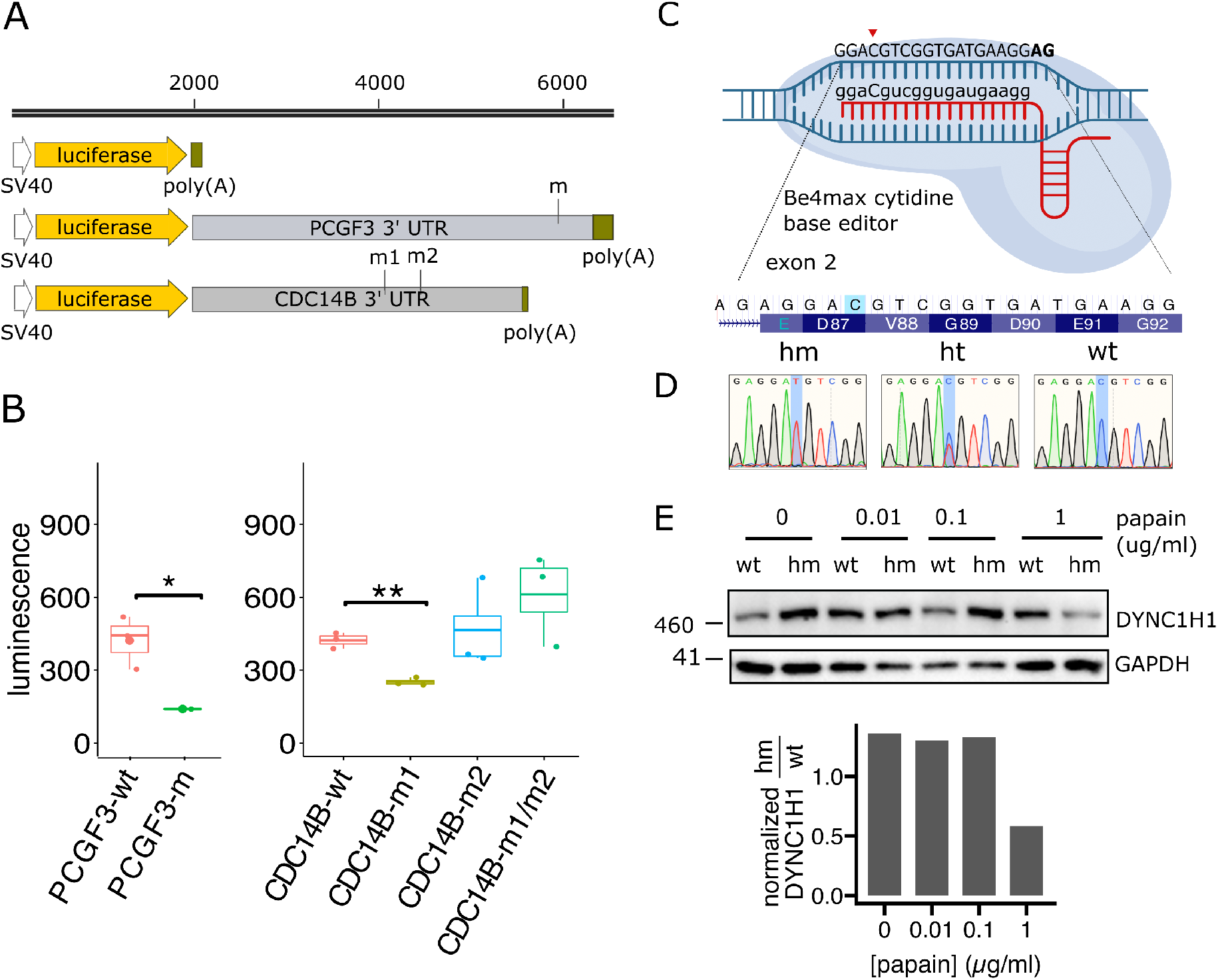
Effects of non-coding and synonymous mutations in *PCGF3, CDC14B* and *DYNC1H1*. A. pGL3-SV40 luciferase reporter plasmids showing *PCGF3* and *CDC14B* 3’ UTRs, additional 3’ sequences with likely poly(A) signals (green) and position of mutations (m, m1, m2). B. Luciferase assays in Y79 cells transfected with the indicated reporter constructs. Significance determined by pair-wise t-test. Box plot lines indicate mean *, p<0.05; **, p<0.01. C. Base-editing strategy for *DYNC1H1* c.261C>T using BE4max cytosine base editor. gRNA sequence shown in lowercase letters except the edited cytidine (arrow). The NG PAM sequence is represented by the bold **AG** on the plus strand. D. Sanger sequence traces of edited homozygous (hm), heterozygous (ht) and wild-type (wt) RB31 clones. E. DYNC1H1 immunoblot of whole cell lysates of homozygous and wild-type CHLA-VC-RB31 clones digested with papain at increasing concentrations. GAPDH is a loading control. Graphs indicate ratio of GAPDH-normalized abundance of DYNC1H1 (hm/wt) for each papain condition.

We next characterized the effects of coding synonymous variant *DYNC1H1* c.261C>T, positioned at the fifth nucleotide of exon 2 and affecting amino acid 261 of 4646 (Fig. 3C). Although *DYNC1H1* is not a widely recognized cancer gene, mutations have been found ^36^. Moreover, the c.261C>T variants’ position is consistent with the increased synonymous variant burden at the 5’ end of cancer gene coding sequences and near the ends of cancer gene internal exons ^35^. We examined the effect of the *DYNC1H1* C.261C>T mutation by generating the variant via cytosine base editing in RB31 cells, chosen for their robust growth (Fig. 3C). After co-transduction with plasmids co-expressing the BE4max cytosine base editor, GFP, and gRNA, transfected GFP+ RB31 cells were isolated by FACS. Bulk GFP+ cultures had ∼ 50% editing rate (Supplementary Fig. S3A) and single cell-derived clones with homozygous, heterozygous, and wild type sequences were identified (Fig. 3D).

Given this variant’s proximity to the exon 2 splice acceptor site, we examined if the mutation impacted splicing via exon-skipping or intron read-through in bulk edited cultures. However, amplification of reverse transcribed cDNA with primer pairs specific to exon 1-2 or exon 1-3 showed no evidence of exon-skipping in the bulk edited or wild-type samples (Supplementary Fig. S3B, C). Likewise, amplification of cDNA intron-exon junction-spanning sequences showed no significant difference, implying there were no changes in retained intron transcripts (Supplementary Fig. S3D). Similarly, we detected no significant differences in *DYNC1H1* RNA abundance as measured by quantitative RT-PCR in *DYNC1H1* regions spanning exons 1-2, exons 5-6, and exons 58-60 (Supplementary Fig. S3E). Thus, we detected no effect of the silent *DYNC1H1* c.261C>T mutation at the RNA level.

As synonymous mutations can alter protein structure, stability, and function via effects on co-translational folding ^35^, we examined DYNC1H1 protein levels and papain protease sensitivity in lysates of *DYNC1H1* mutant and wild type clones. Whereas homozygous mutant DYNC1H1 protein levels were reproducibly higher than wild type DYNC1H1 in cell lysates with no, 0.01 μg/ml, or 0.1 μg/ml papain, they were lower in lysates treated with 1 μg/ml papain (Fig. 3E). Thus, the *DYNC1H1* c.261C>T coding-silent mutation increased sensitivity to protease digestion.

### An acquired biological process during cell line establishment

To assess whether recurrently altered genomic changes and biological process ontologies are selected in retinoblastoma-derived cell lines, we performed WES on 12 tumor-matched retinoblastoma cell lines sampled during exponential growth after a total of 100-200 days in culture (Fig. 4A, Supplementary Table S5). Exome based analysis of DNA copy number changes revealed one or more retinoblastoma SCNA (1q+, 2p+, 6p+, 7q+, 16q-, and 19q+) in all CHLA-VC-RB tumors and focal *BCOR* deletions in RB14 and RB31 (Fig. 4B, Supplementary Table S6). Most SCNAs were subclonal when compared to *RB1* variant frequency in the same primary tumors, and most cell lines were enriched for retinoblastoma SCNAs relative to the parental tumor (Fig. 4B). For example, tumor CHLA-VC-RB20 had subclonal 16q loss while the cell line had clonal 16q loss, enhanced 1q gain, and *de novo* 2p gain. Tumor CHLA-VC-RB49 had subclonal 1q+, 2p+, 6p+, and 16q-while these SCNAs were clonal in the cell line. As an exception, 16q-was detected in tumor CHLA-VC-RB46 but not in the corresponding cell line. CN-LOH was largely concordant with SCNAs (Fig. 4B-C and Supplementary Tables S6-S7).

**Figure 4.**
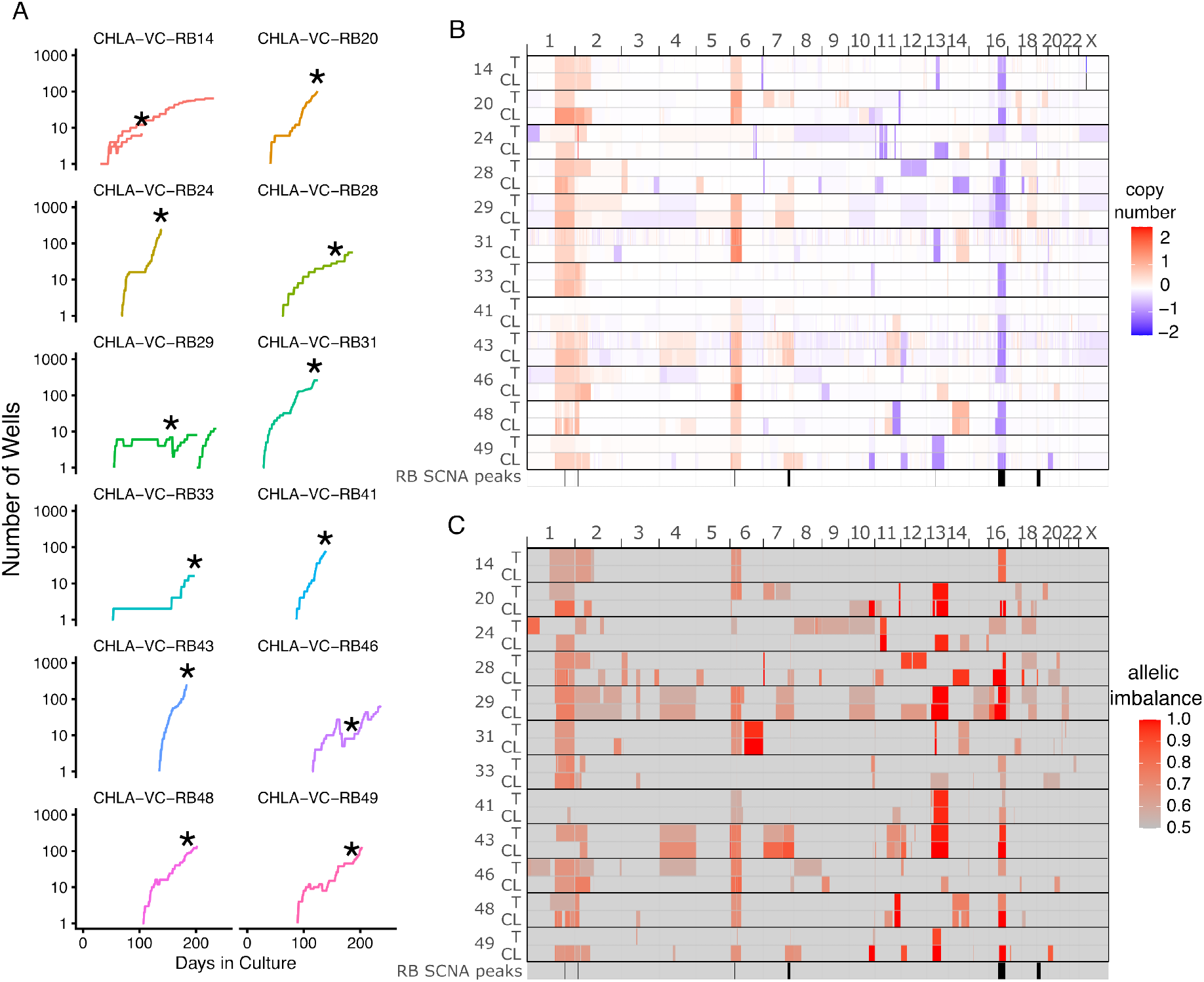
DNA copy number and allelic imbalance changes in CHLA-VC-RB cell lines. A. CHLA-VC-RB cell growth. X-axis represents cumulative days in culture post-explant. Y-axis represents 96-well equivalents after initial seeding with ∼ 1-5×10^6^ cells. Growth was defined by culture well numbers rather than cell numbers to minimize cell dissociation and maximize viability. Asterisks, time point used for exome sequencing. Cultures with stalled growth were subsequently re-expanded in culture (CHLA-VC-RB14) or xenograft (CHLA-VC-RB29). B. Somatic copy number alterations (SCNAs) in tumor (T) - cell line (CL) pairs and represented on log_2_ scale. Blue; loss; red; gain. C. Allelic imbalance heat map representing mirrored B allele frequency in tumor – cell line pairs. RB SCNA peaks indicated below each panel represent the most recurrent SCNA regions ^**7**^.

We also detected differences in the variant spectrum and VAFs in matched tumor and cell line somatic mutations (Fig. 5). We detected 59 SNVs in the 12 CHLA-VC-RB cell lines (mean: 4.92; median: 4.5) (Fig. 1A, B). 31 of 60 (52%) variants detected in tumors were retained in cell lines and 31 of 59 (53%) mutations detected in cell lines had detectable antecedents in their primary tumors. Among genes altered in at least two retinoblastomas, *PAN2* G793A VAF increased from 0.3 to 0.51 (p<0.01) and *SAMD9* P63S VAF increased from 0.02 to 0.08 (p<0.001) during CL29 and CL43 establishment (Fig. 5A). In contrast, *NAF1* R255Q VAF decreased from 0.14 to undetected in CL24 (p<0.01), *BCOR* R1400IfsX12 declined from 0.21 to undetected in CL28 (p < 0.0001), and *BCOR* G756X declined albeit not significantly from 0.29 to 0.20 in CL33 (Fig. 5B). Thus, the recurrently altered *BCOR* and *NAF1* were not selected during cell line establishment.

**Figure 5.**
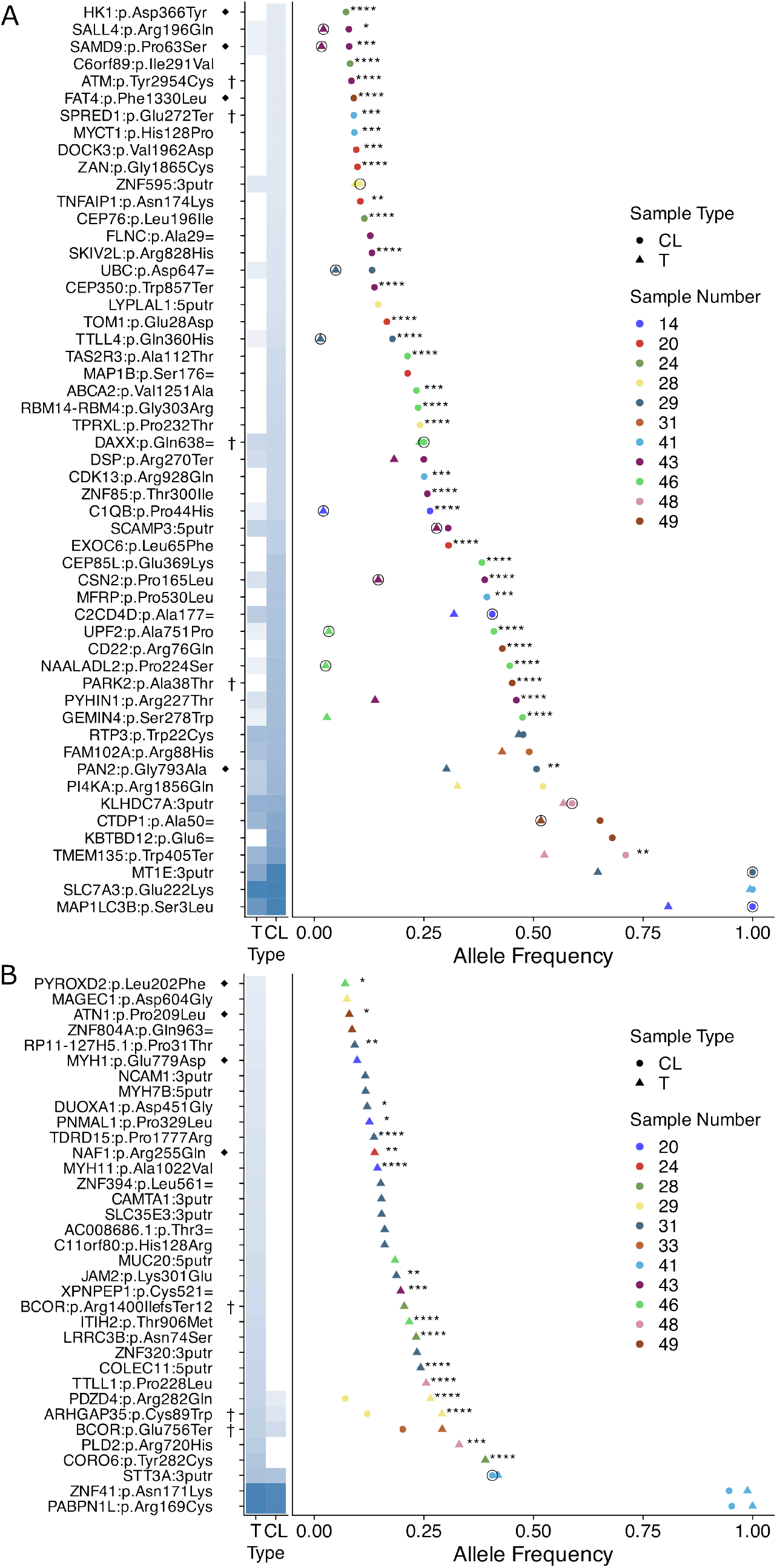
Curated somatic variants in CHLA-VC-RB tumor and cell lines. Variants in tumors (triangles) and cell lines (circles) are displayed in ascending VAF order for samples with increased (A) or decreased (B) variant frequency in tumor-derived cell lines, as shown in heat maps at left. Circled icons indicate variants initially detected in a cell line and subsequently detected exclusively in the matched tumor. ◆, alleles confirmed by targeted sequencing. †, genes listed in targeted sequencing panels (UCSF500 or MSK-IMPACT). Asterisks, significantly altered allele frequencies in tumor-cell line pairs (*, p<0.05; **, p<0.01; ***, p<0.001; ****, p<0.0001; Fisher exact test).

We further asked if CHLA-VC-RB cell lines acquired mutations in biological processes distinct from those in primary tumors. Over-representation analyses of the 59 somatic variants in 12 CHLA-VC-RB cell lines revealed enrichment of only three ontologies with FDR < 0.25, all relating to positive regulation of DNA damage response by p53 (Fig. 2B). Each ontology was specified by non-synonymous variants of *ATM, SPRED1*, and *PYHIN1* ^37,38^, which had higher VAFs in cell culture then in their originating tumors (Fig. 5A). In contrast, p53-related ontologies were not over-represented among somatic variants in 168 retinoblastoma tumors (Supplementary Table S11), implying that altered p53 signaling is selected in at least some retinoblastoma cell lines.

## Discussion

This study aimed to assess contributions of non-synonymous, synonymous, and non-coding genomic abnormalities to recurrently altered cellular processes during retinoblastoma progression. Retinoblastoma is an apt setting to study such contributions as the low mutation rate and early age of onset likely limits the number of passenger mutations.

Combining variants in 168 tumors revealed over-representation of previously unrecognized progression-related processes. The most significantly over-represented ontology was *histone monoubiquitination*, reflecting the over-arching impact of *BCOR*, a component of non-canonical Polycomb repressive complex 1 (PRC1) that mediates histone H2A monoubiquitination ^39^ and a recognized recurrent secondary mutation in retinoblastoma and other cancers ^13,40^. As a more novel finding, we detected over-representation of variants related to assembly of the CCR4/NOT1 complex and mRNA P body (*CNOT1, CNOT2, PAN2*, and *DYNC1H1*), which mediate mRNA deadenylation and degradation ^41,42^. The *CNOT2* variant (c.162insA) was an inactivating frame-shift mutation whereas the *PAN2* variants G793A and D241H (Supplementary Table S1) were in the C-terminal hydrolase and linker regions that bind PAN3 ^43^, suggesting that each mutant affects PAN2-PAN3 interactions and mRNA deadenylation. Notably, in *Drosophila*, knockdown of different CNOT components increased formation of Rb family-deficient retinal tumors ^44^. Thus, our data implicate the disruption of CCR4/NOT1 complexes in human as well as in *Drosophila* pRB-deficient photoreceptor cancers. Other over-represented biological process ontologies also included *mitotic cytokinesis* and *sister chromatid segregation, cell adhesion mediated by integrin, semaphorin-plexin signaling*, and *phosphatidylinositol phosphorylation*, suggesting additional tumor progression-related mechanisms. Using pathway analysis, Field *et al*. observed that some secondary mutations could be connected in a protein interaction network that included the estrogen-related receptor ESRRG, which in turn was implicated in the retinoblastoma cell hypoxic response ^14^. ESRRG- and hypoxic response-related ontologies were not enriched in our over-representation analyses or in an Ingenuity pathway analysis of the 542 somatic variants, though this could relate to deficiencies in ontology enrichment databases.

Strikingly, most of the over-represented ontologies were more significantly over-represented when synonymous coding and non-coding UTR mutations were included (Fig. 2A). While often considered passenger mutations, synonymous and UTR variants may act via RNA sequence effects on splicing, translation start-site selection, translation efficiency, and co-translational folding that alter protein function ^34^. Moreover, they are far more likely to impact oncogenesis in cancers, such as retinoblastoma, with low mutational load and thus fewer passenger mutations ^35^. Indeed, nonsynonymous as well as synonymous and non-coding variants had a similar propensity to contribute to over-represented biological process ontologies and three out of three such variants examined had deleterious effects.

Specifically, a 3’ UTR variant of *PCGF3* significantly reduced gene expression. As *PCGF3* encodes a protein that recruits BCOR to Polycomb repressor complex 1 ^45^, its reduced expression is expected to impair BCOR’s histone monoubiquitination role in a manner similar to *BCOR* loss. Of two 3’ UTR variants in *CDC14B*, the m1 mutation impaired *CDC14B* expression whereas m2 reversed this effect. Thus, as a possible scenario, an m1-induced *CDC14B* down-regulation might subvert the *CDC14B* role in the spindle checkpoint to enable tumor evolution ^46^, while m2 could restore *CDC14B* expression to ensure optimal growth of the evolved tumor. Finally, we found that a synonymous *DYNC1H1* c.261C>T variant sensitized DYNC1H1 to protease digestion, evidence of altered protein structure. Interestingly, *DYNC1H1* is a long (19.9 kb) mRNA that accumulates in translation factories ^47^ and is enriched in stress granules in response to translation stressors ^41^, consistent with its having a regulated co-translational folding mechanism. Moreover, the *DYNC1H1* c.261C>T mutation has properties typical of synonymous mutations in cancer genes, including its position in the first 10% of the coding sequences near an intron-exon border ^35^, which can affect co-translational folding ^48^. More routine analysis of synonymous and non-coding mutations is likely to provide further insight into retinoblastoma progression mechanisms.

This study also characterized the relationship of retinoblastoma cell lines to their antecedent tumors. Genomically well-characterized cell lines are needed to dissect the role of cancer progression-related genomic changes in their native genomic backgrounds. However, tumor heterogeneity necessitates that some subclonal variants are not retained in cell lines, whereas ongoing mutagenesis and clonal selection ensures that cell lines have novel variants not detected in the tumors (Fig. 5). Concordantly, we found that only ∼ 50% of SNVs were enriched during cell line establishment, and that even repeatedly altered genes implicated in retinoblastoma oncogenesis were not always retained. Of particular note, two inactivating (premature termination) *BCOR* variant alleles were depleted (Fig. 5), implying that *BCOR* variants that are selected in tumors need not be selected during cell line establishment. In line with this finding, Field *et al*. have proposed that *BCOR* represses *ESRRG1* to enable a hypoxic survival response, a function that would have no advantage in the hyperoxic tissue culture setting ^14^. CHLA-VC-RB cell lines also had over-representation of *ATM, SPRED1*, and *PYHIN1* variants related to the p53-mediated DNA damage response. Notably, the *SPRED1* variant appeared in CHLA-VC-RB41, which was the only cell line that lacked gain of the chromosome 1q region harboring *MDM4* and implicated in impairing p53 function ^49^. Thus, it will be important to assess whether p53 signaling may be impaired by multiple mechanisms during cell line establishment.

In sum, these studies reveal that retinoblastoma progression is associated with accumulation of subclonal gene variants that affect diverse cell signaling pathways. While over-representation of the *BCOR*-driven histone monoubiquitination term was expected, we also detected over-representation of variants that affect assembly of RNP complexes that regulate mRNA stability and have been implicated in retinoblastoma genesis in a *Drosophila* model. Most of the affected biological process ontologies were more significantly over-represented when including non-coding UTR variants and synonymous variants that could affect gene function at the RNA or protein structure level. Finally, our data suggest that the effects of some progression-related variants may be challenging to decipher in cell lines which lack important *in vivo* constraints and may be selected to impair the p53-mediated DNA damage response.

## Ethics approval statement

Retinoblastoma tumors and normal tissues were obtained under protocols approved by the Children’s Hospital Los Angeles and Children’s Oncology Group (COG) institutional review boards.

## Patient consent statement

Guardians of all patients provided informed consent for donation of tumor for research purposes.

## Permission to reproduce material from other sources

Not applicable

## Clinical trial registration

Not applicable

## Acknowledgments

We thank Carly Stewart and Heather Davidson for managing cell lines, Irsan Kooi for scripts, Irsan Kooi and Martin Triska for helpful discussions, and core facilities at The Saban Research Institute of Children’s Hospital Los Angeles: the Flow Cytometry (FACS) Core for cell sorting assistance and Stem Cell Analytics Core for transfection assistance. This study was supported by The Nautica Malibu Triathlon produced by MSP Inc (DC), The St. Baldrick’s Foundation (DC), NIH K08CA232344 (JLB), NIH K08EY030924 (AN), the Las Madrinas Endowment in Experimental Therapeutics for Ophthalmology (AN), a Research to Prevent Blindness Career Development Award (AN), an unrestricted grant to the USC Department of Ophthalmology from Research to Prevent Blindness, The Larry and Celia Moh Foundation, The Neonatal Blindness Research Fund, The Knights Templar Eye Foundation (AN and DC), and NIH R01CA137124 (DC).

## Supplementary Data

### Supplementary Figures

**Figure S1**. Exome sequencing coverage by sample type.

**Figure S2**. Expression and conservation of *PCGF3* and *CDC14B* variant sequences.

**Figure S3**. *DYNC1H1* splicing and RNA expression not impacted by C.261C>T.

**Figure S4**. DNA copy number, allelic imbalance, and growth of CHLA-RB cell lines.

### Supplementary Tables

**Table S1**. Somatic exome mutations beyond *RB1* in fully reported retinoblastoma exome, genome, and targeted sequencing studies.

**Table S2**. Sample coverage and variants beyond RB1 in retinoblastoma sequencing studies.

**Table S3**. STR typing of CHLA-VC-RB and CHLA-RB samples.

**Table S4**. Oligonucleotide primers used for PCR and site-directed mutagenesis.

**Table S5**. Patients in this and prior retinoblastoma sequencing studies.

**Table S6**. Copy number alterations in CHLA-VC-RB and CHLA-RB tumor and cell lines.

**Table S7**. Allelic imbalance in CHLA-VC-RB and CHLA-RB tumors and cell lines.

**Table S8**. Somatic variants detected in CHLA-VC-RB tumors and cell lines.

**Table S9**. Biological process ontologies of somatically altered genes in sequenced retinoblastoma tumors (n=160) and cell lines (n=12).

Table S10. Enrichment analysis iterations assessing synonymous and UTR variant contribution.

**Table S11**. Somatic variants detected in CHLA-RB cell lines.

**Figure S1.**
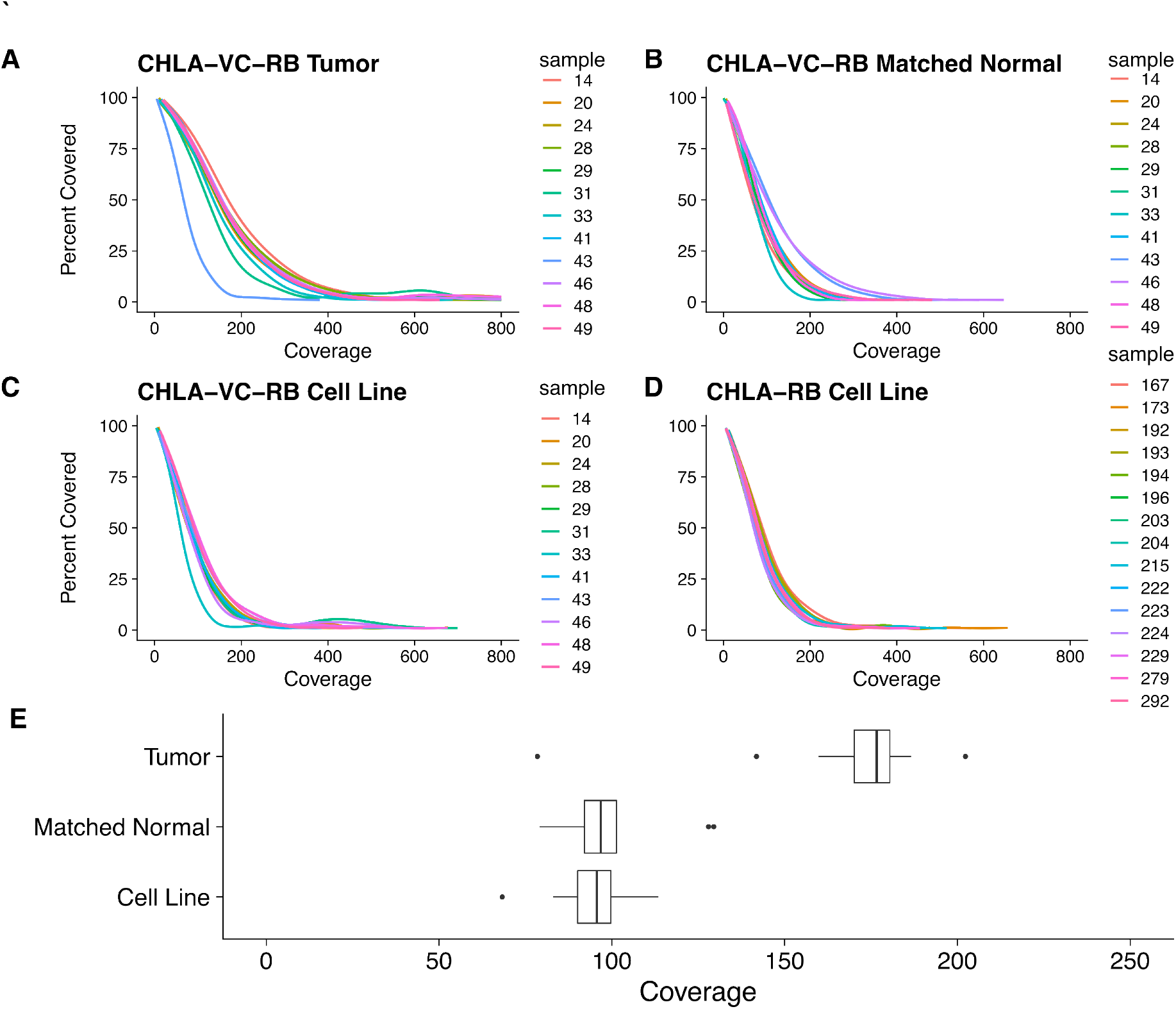
Exome sequencing coverage by sample type. (A-D); Percent of exome covered at read depth for CHLA-VC-RB tumors (A); CHLA VC-RB matched normal DNAs (B); CHLA-VC-RB cell lines (C); and CHLA-RB cell lines (D). (E); Box plot representing mean sequencing coverage of CHLA-VC-RB samples.

**Figure S2.**
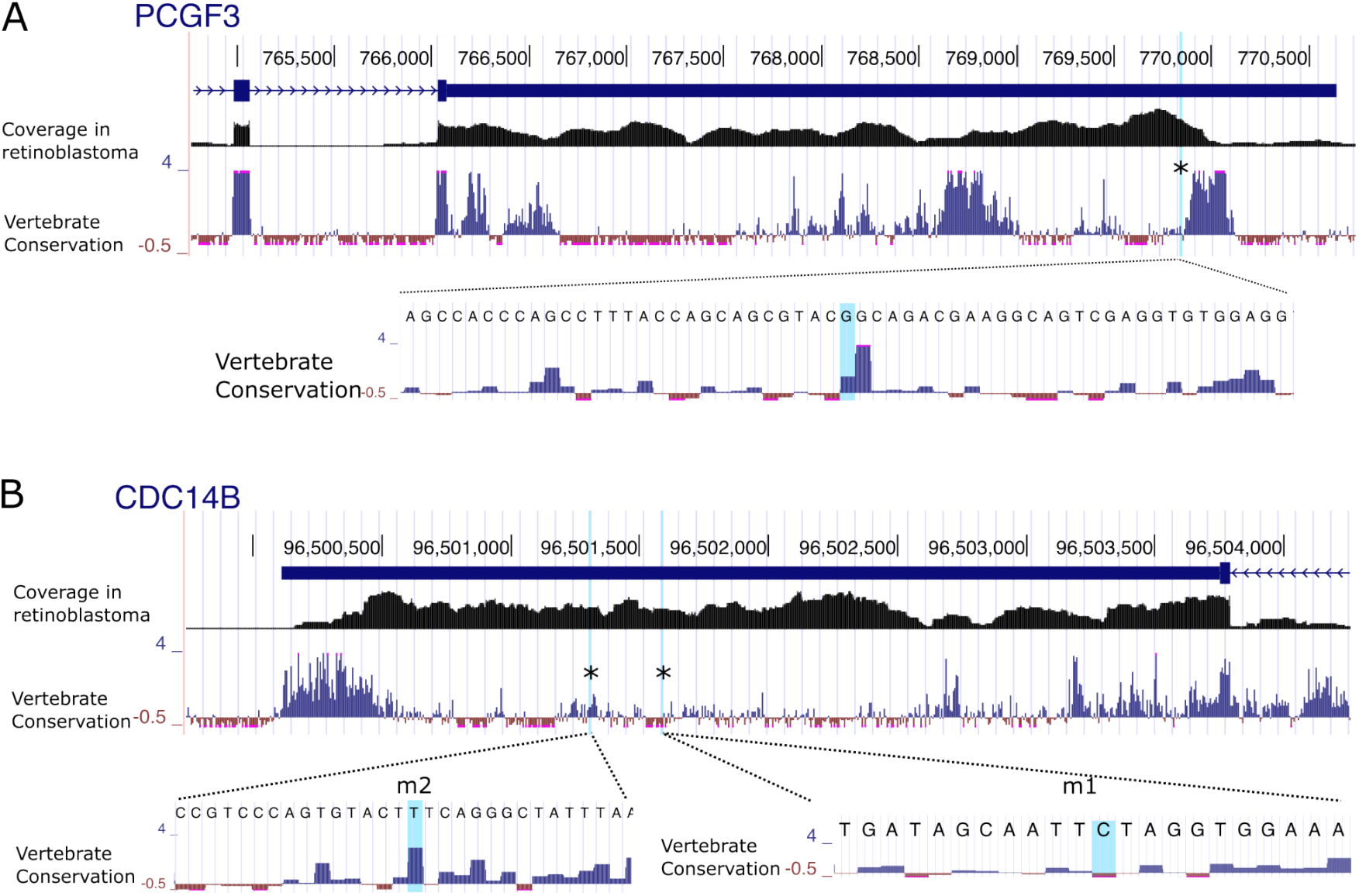
Expression and conservation of *PCGF3* and *CDC14B* variant sequences. UCSC Browser views of *PCGF3* (A) and *CDC14B* (B) 3’ UTRs, displaying RNAseq expression in a representative retinoblastoma tumor (GSE125903) and vertebrate conservation at low and high resolution. Note genomic *CDC14B* sequences represent the cDNA *minus* strand. Asterisks and blue shading indicate variant positions.

**Figure S3.**
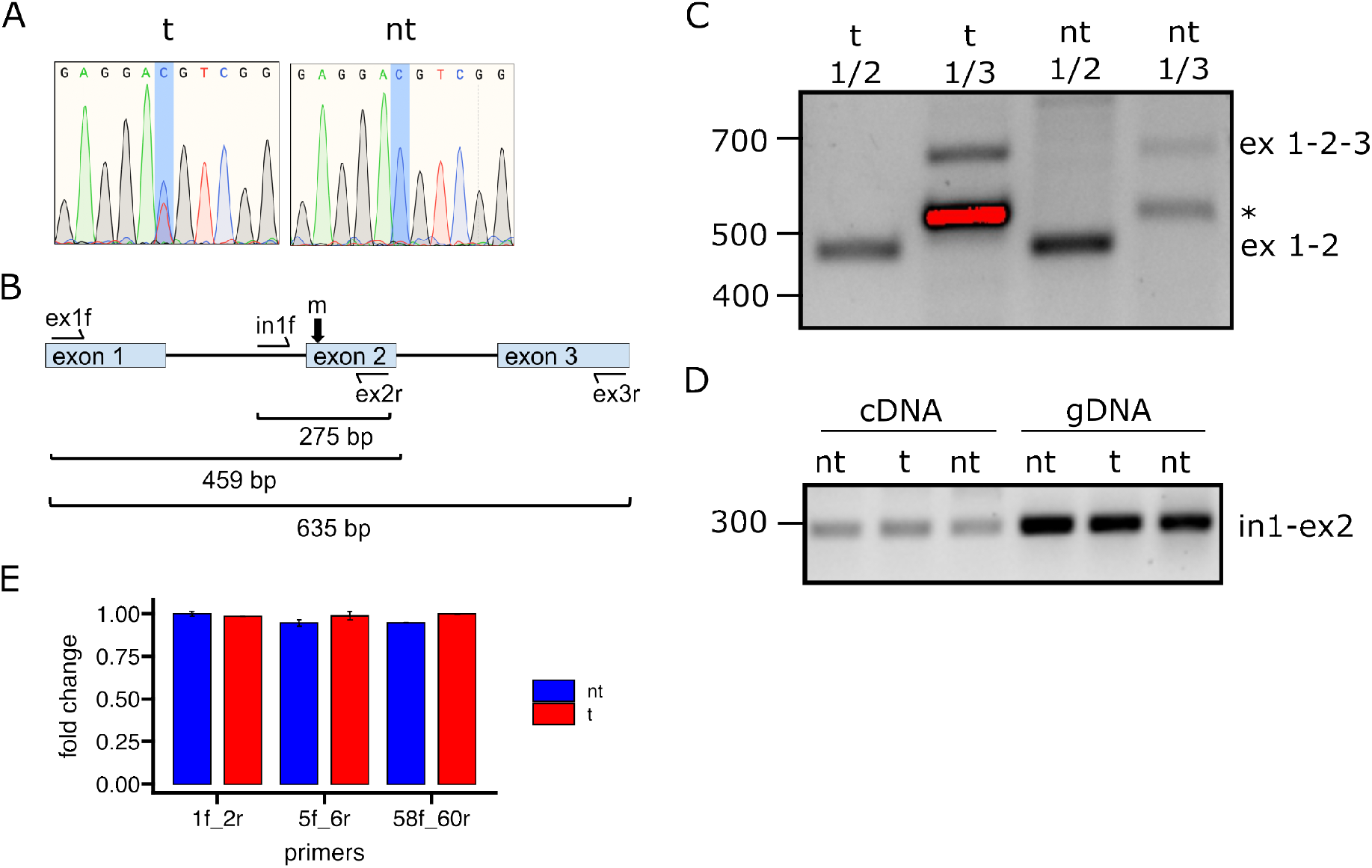
*DYNC1H1* splicing and RNA expression not impacted by C.261C>T. A. Sanger sequence traces of base-edited *DYNC1H1* region at 24 h after co-transfection of CHLA-VC-RB31 cells with dCas9-cytosine base editor and either targeting (t) or non-targeting (nt) gRNAs, followed by isolation of transfected GFP+ cells, showing ∼ 50% C>T conversion at the targeted cytidine. B. *DYNC1H1* exons 1,2,3 with PCR primer positions and predicted product sizes. C. RT-PCR analysis of RNA from base-edited (t) or non-targeted (nt) bulk RB31 cultures using *DYNC1H1* ex1f/ex2r and ex1f/ex3r primers. Positions of sequence-verified exon 1-2 and exon 1-2-3 products are indicated. An additional band in ex1f/ex3r reactions (*) was non-specific and did not contain exon 1-3 (exon 2-skipped) product in nested PCR analyses. D. Amplification of cDNA (generated with DNase-treated RNA) and gDNA products using intron1f/exon2r primer pairs E. RT-qPCR analysis of *DYNC1H1* expression in targeted (C.261C>T) or one non-targeted bulk culture using primers amplifying exons 1-2, exons 5-6, and exons 58-60. Error bars, standard deviation of duplicate qPCR reactions.

**Figure S4.**
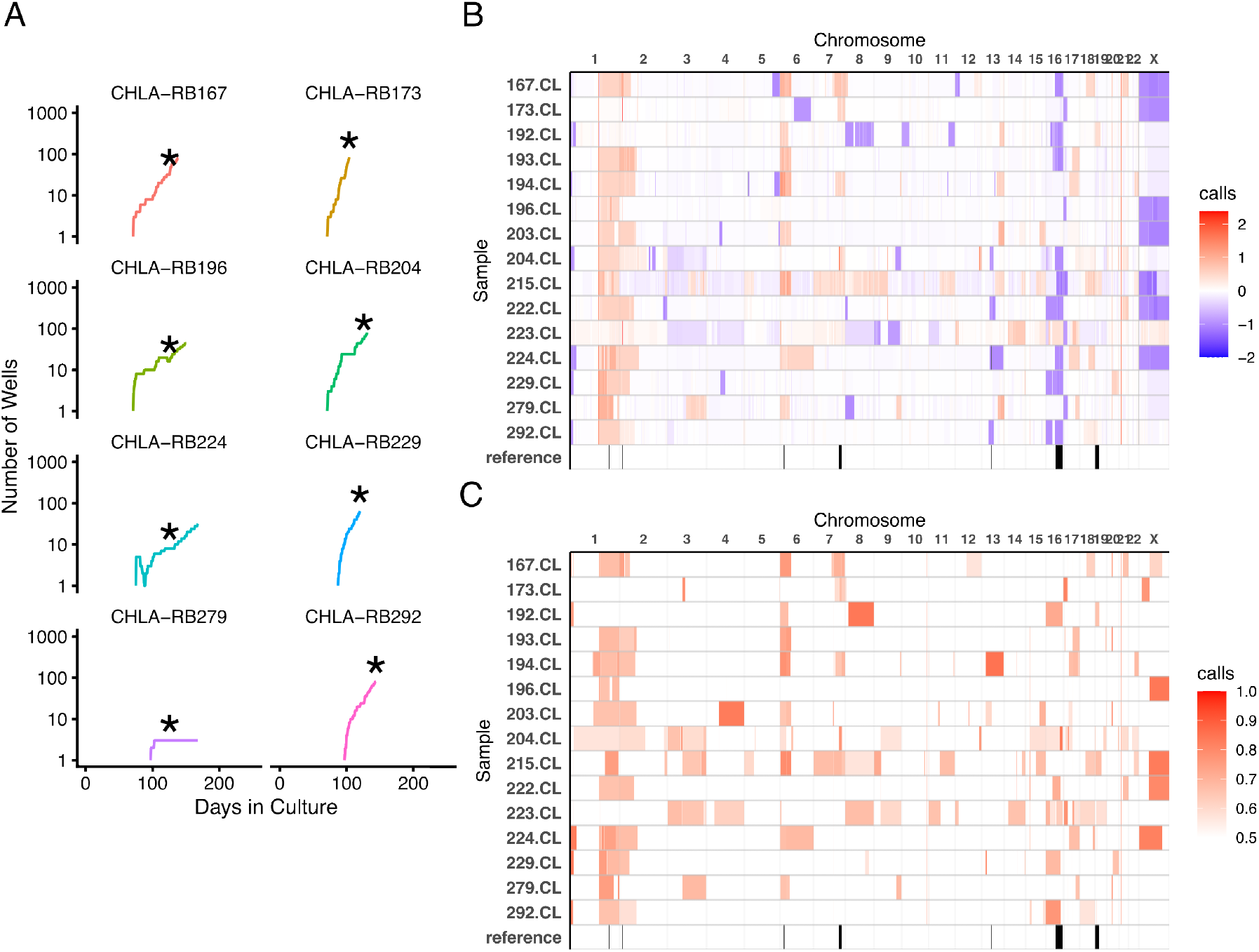
CHLA-RB cell line growth, DNA copy number, and allelic imbalance. A. Growth of eight CHLA-RB cell lines. X-axis represents cumulative days in culture post-explant. Y-axis represents 96-well equivalents after initial seeding with ∼ 1-5×10^6^ cells. Growth was defined by culture well numbers rather than cell numbers to minimize cell dissociation and maximize viability. Asterisks, time point used for exome sequencing. B. SCNAs defined by copywriteR represented on log_2_ scale. Blue; loss; red; gain. C. Allelic imbalance heat map representing mirrored B allele frequency. RB SCNA peaks indicated below each panel represent the most recurrent SCNA regions ^**7**^.

**Table S4.**
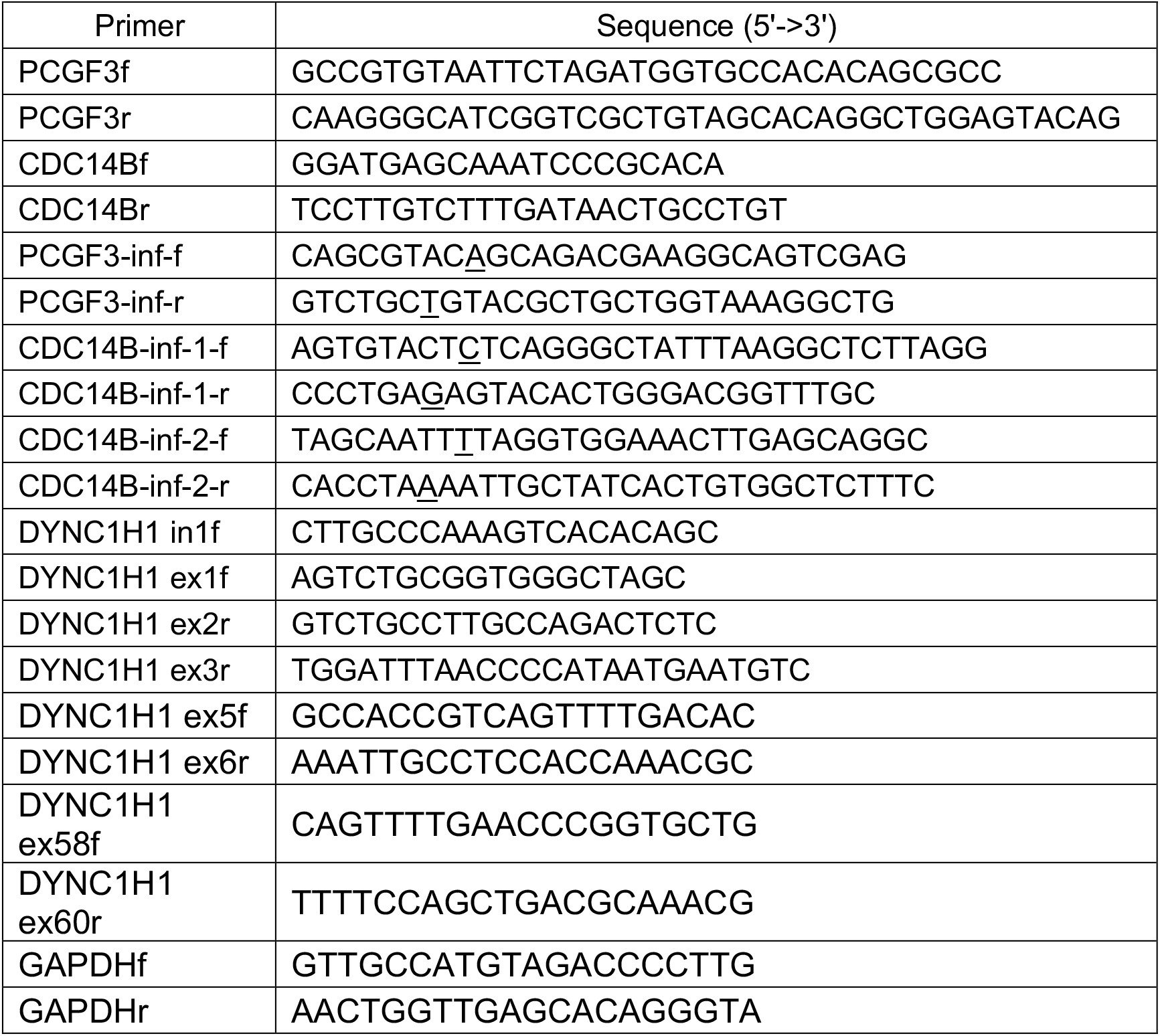
Oligonucleotide primers used for PCR and site-directed mutagenesis. Underlined bases in In-Fusion mutagenesis (inf) primers represent 3’ UTR mutations.

